# The *Thermosynechococcus* genus: wide environmental distribution, but a highly conserved genomic core

**DOI:** 10.1101/2020.10.20.346296

**Authors:** A. Paulina Prondzinsky, Sarah J. Berkemer, Lewis M. Ward, Shawn E. McGlynn

**Affiliations:** Department of Chemical Science and Engineering, Tokyo Institute of Technology, Ookayama, Meguro-ku, Tokyo, Japan; Earth-Life Science Institute, Tokyo Institute of Technology, Ookayama, Meguro-ku, Tokyo, Japan; Bioinformatics Group, Department of Computer Science, University Leipzig, Leipzig, Germany; Competence Center for Scalable Data Services and Solutions, Dresden/Leipzig, Germany; Department of Earth and Planetary Sciences, Harvard University, Cambridge, MA, USA

**Keywords:** cyanobacteria, great oxygenation event, hot springs, comparative genomics, pan-genome, *Thermosynechococcus*

## Abstract

Cyanobacteria thrive in very diverse environments. However, questions remain about possible growth limitations in ancient environmental conditions. As a single genus, the *Thermosynechococcus* are cosmopolitan and live in chemically diverse habitats. To understand the genetic basis for this, we compared the protein coding component of *Thermosynechococcus* genomes. Supplementing the known genetic diversity of *Thermosynechococcus*, we report draft metagenome-assembled genomes of two *Thermosynechococcus* recovered from ferrous carbonate hot springs in Japan. We find that as a genus, *Thermosynechococcus* is genomically conserved, having a small pan-genome with few accessory genes per individual strain and only 14 putative orthologous protein groups appearing in all *Thermosynechococcus* but not in any other cyanobacteria in our analysis. Furthermore, by comparing orthologous protein groups, including an analysis of genes encoding proteins with an iron related function (uptake, storage or utilization), no clear differences in genetic content, or adaptive mechanisms could be detected between genus members, despite the range of environments they inhabit. Overall, our results highlight a seemingly innate ability for *Thermosynechococcus* to inhabit diverse habitats without having undergone substantial genomic adaptation to accommodate this. The finding of *Thermosynechococcus* in both hot and high iron environments without adaptation recognizable from the perspective of the proteome has implications for understanding the basis of thermophily within this clade, and also for understanding the possible genetic basis for high iron tolerance in cyanobacteria on early Earth. The conserved core genome may be indicative of an allopatric lifestyle – or reduced genetic complexity of hot spring habitats relative to other environments.

## Introduction

Water oxidizing cyanobacteria fundamentally altered the distribution of carbon and electrons on Earth (Canfield, 1998; Raven, 2009; Ward and Shih, 2019). A marker of this reorganization is in the Great Oxygenation Event (GOE), which marks a major transition in the evolution of life on Earth ~2.3 billion years ago (Lyons *et al.*, 2014; Fischer *et al.*, 2016). While the GOE is widely accepted to have been driven by the production of O_2_ by oxygenic photosynthesis performed by members of the cyanobacteria, the timing and proximal trigger for the GOE is debated. At least six hypotheses for the timing of the GOE have been considered i) the evolution of oxygenic photosynthesis by cyanobacteria just before the GOE (Fischer *et al.*, 2016; Shih *et al.*, 2017), ii) an earlier evolution of cyanobacteria with O_2_ accumulation delayed due to the transition of cyanobacteria from small-scale freshwater to large-scale marine environments (Sánchez-Baracaldo, 2015), iii) the transition from unicellular to multicellular organisms for increased evolutionary success (Schirrmeister *et al.*, 2015), iv) the inhibition of early cyanobacteria due to high iron concentrations (Swanner, Mloszewska, *et al.*, 2015; Swanner, Wu, *et al.*, 2015), v) a possible nitrogen throttle on cyanobacterial growth (Fennel *et al.*, 2005; Shi and Falkowski, 2008; Allen *et al.*, 2019), vi) or depressed Archaean productivity due to phosphate availability (Hao *et al.* 2020).

When and where the last common ancestor of cyanobacteria emerged is a matter of debate (Sánchez-Baracaldo, 2015; Shih *et al.*, 2017a; Tria *et al.*, 2017) and cyanobacterial taxonomy continues to be refined (Knoll 2006; Tomitani *et al.*, 2006; Shih *et al.*, 2013; Soo *et al.*, 2019; Parks *et al.*, 2020). Although uncertainty exists as to how much can be learned about past ecology from contemporary biology, an increased understanding of today’s organisms may help us generate, or reject hypotheses about former evolutionary states.

Cyanobacteria are found in a wide range of environments – the knowledge of which continues to expand (Whitton, 1992; Puente-Sánchez *et al.*, 2018; Callieri *et al.*, 2019). As a single phylogenetic group, members of the *Thermosynechococcus* genus have been documented to inhabit a range of chemical environments, some of which can be seen as analogous to the Proterozoic environments on the early Earth (Ward *et al.*, 2019).

Phylogenetic analysis of the *Thermosynechococcus* indicates they are a relatively recent divergence (Sánchez-Baracaldo, 2015; Shih *et al.*, 2017). As a coherent group which spans a limited evolutionary distance, they provide a test case for querying how much environmental adaptation can be achieved within a limited time frame; that *Thermosynechococcus* inhabit a wide array of environments implies that rapid adaptation is possible, but the basis for this is unknown. Those *Thermosynechococcus* with genomes available are from hot springs which vary in temperature from 44 – 94 °C, pH ranging from 5.4 – 9.3, sulfate concentrations between 0.06 mM and 17.4 mM and iron concentrations between 0.4 μM and 261 μM (Table 1) at their source (however it must be emphasized that the source waters are not where the organisms reside unless specifically noted and the source water chemistry only gives a starting point of a gradient).

**Table 1:** Geochemical parameters for hot spring source waters. Values in parentheses indicate the geochemical values of the sites where *Thermosynechococcus* sequences were observed, if known. Other values indicate the source water geochemistry of each spring, which can be used as a reference point for the start of a gradient in cases in which the explicit site where *Thermosynechococcus* sequences were observed is unknown. ^a^Concentrations were derived from geochemical modelling in Mei-hua *et al.* (2015). ^b^Iron for Jinata and Okuoku-Hachikurou hot springs is ferrous, all others are totals of ferrous plus ferric iron. References are related to the first publication of the strains or the geochemistry of the hot spring. *information available online through local governments, last accessed in 11/2019.

| Name | Strain name<br>(Thermosynecho<br>coccus) | Max.<br>Temperat<br>ure [°C] | pH | Sulfate<br>[mM] | Total<br>Iron<br>[μM] | References |
| --- | --- | --- | --- | --- | --- | --- |
| Kangding<br>Geotherm<br>al Field<br>Lianhua<br>Lake<br>hotspring<br>(Sichuan,<br>China) | *T. elongatus*<br>PKUAC-SCTE542<br>(*elongatus*<br>PKUAC) | 94<br>(67.2) | 6.35 –<br>8.84<br>(7.95) | 1.2 | 10.4<sup>a</sup> | Mei-hua *et al.*,<br>2015; Guo *et al.*,<br>2017; Tang *et al.*, 2018 |
| Okuoku-hachikurou<br>Hot Spring<br>(Akita,<br>Japan) | *Ca. T.*<br>*nakabusensis*<br>OHK43 (OHK) | 44 | 6.8<br>(6.8) | 6.5 | 114<sup>b</sup><br>(> 100) | Ward *et al.*,<br>2017 |
| Yunomine<br>Hot Spring<br>(Wakayam<br>a, Japan) | *T. vulcanus*<br>NIES2134/1-2178 (*vulcanus*) | 91 | 8 | 0.06 –<br>0.229 | < 1.8 | Ichikuni and<br>Kikuchi, 1972<br>Onsen<br>information<br>sheet* |
| Nakabusa<br>Hot Spring<br>(Nagano,<br>Japan) | *T. nakabusensis*<br>NK55a<br>(NK55a) | 76 | 8.5 –<br>9 | 0.218 –<br>0.246 | 0.4 | Nakagawa and<br>Fukui, 2002;<br>Nakamura *et al.*,<br>2002; Stolyar *et al.*, 2014<br>Onsen<br>information<br>sheet* |
| Jinata Hot<br>Spring<br>(Shikinejim<br>a, Tokyo,<br>Japan) | *Ca. T.*<br>*nakabusensis*<br>Jinata (J003)<br>*T. sp.*<br>M3746_W2019_0<br>13 (Jinata 2) | 63<br>(37 – 46) | 5.4<br>(6.7) | 17.4 | 261<sup>b</sup><br>(> 100) | Ward *et al.*,<br>2019, Alcorta *et al.*, 2020 |
| Chung-Lun Hot Spring (Taiwan) | *T. sp.* CL1/1-2178 (CL1) | 62 | 9.3 | 1.35 – 1.39 | 0.6 | Yeh *et al.*, 2005; Hsueh *et al.*, 2007; Maity *et al.*, 2011; Cheng *et al.*, 2020 |
| Beppu Hot Spring Kamegawa Shinoyu (Oita, Japan) | *T. elongatus* BP1/1-2178 (*elongatus* BP1) | 78 | 6.8 | 1.09 | 3.6 | Onsen information sheet* |
| Shivlinga hot spring, Ladhak, India | *T. sp.* M46_R2017_013 (Shivlinga)<br>*excluded in genome comparison due to low completeness | 70 (46) | 7 (8) | 1 | No data | Alcorta *et al.*, 2020; Roy *et al.*, 2020 |
| Tattapani, India | *T. sp.* M55_K2018_012 (Tattapani 1)<br>*T. sp.* M98_K2018_005 (Tattapani 2) | 98 (55) | 7.7 (7.9) | No data | 0.49 | Kaushal *et al.*, 2018; Alcorta *et al.*, 2020; Saxena *et al.*, 2017 |

The *Ca. T. nakabusensis* OHK43 (OHK43) genome was recovered from material at the edge of the Okuoku-hachikurou Onsen (OHK) source pool where iron concentrations were measured to be 114μM, and the *Ca. T. nakabusensis* Jinata (J003) genome was recovered at Jinata hot spring along a gradient where iron concentrations were 100 μM and greater (Ward et al 2017 and Ward et al 2019). In contrast, the other springs where *Thermosynechococcus* genomes originate are from hot springs with iron concentrations below the suggested toxicity threshold of tens to hundreds of micromolar Fe(II) suggested previously (Swanner, Mloszewska, *et al.* 2015). Furthermore, Nakabusa hot spring is sulfidic at 0.1 mM sulfide, while sulfide concentrations for Jinata and OHK were below the detection limit of a Cline assay (unpublished data), showing additional differences in geochemistry, which together are shown in Table 1.

Seminal work by Papke *et al.* (2003) on phylotype:geographical relationships of *Thermosynechococcus* posed questions as to how these organisms could be so widely distributed: in their analysis, the distribution of *Thermosynechococcus* could not be explained by measured geochemical parameters from their respective environments of origin. *Thermosynechococcus* thus appear to be cosmopolitan, but the basis for this remains unresolved. Motivated by the finding of *Thermosynechococcus* members in ferrous iron carbonate hot springs that we have been studying (Ward *et al.*, 2017, 2019), we sought to test the hypothesis that *Thermosynechococcus* members in high iron springs may have genetically resolvable traits associated with iron adaptation, by comparing to strains which are from lower iron environments.

## Methods

### Genome recovery

The OHK43 genome was recovered from genome-resolved metagenomic sequencing of samples from Okuoku-hachikurou Onsen (OHK) in Akita Prefecture, Japan, following methods described previously (Ward *et al.*, 2019, 2020) and described briefly here. Samples of thin biofilms in the outflow of the hot spring were sampled for metagenomic sequencing in September 2016. DNA was preserved in the field with a Zymo Terralyzer BashingBead Matrix and Xpedition Lysis Buffer (Zymo Research, Irvine, CA) after disruption of cells in polyethylene sample tubes via attachment to and operation of a cordless reciprocating saw (Makita JR101DZ). Microbial DNA was extracted and purified after return to the lab with a Zymo Soil/Fecal DNA extraction kit (Zymo Research, Irvine, CA). Quantification of DNA was performed with a Qubit 3.0 fluorimeter (Life Technologies, Carlsbad, CA). DNA was submitted to SeqMatic LLC (Fremont, CA) for library preparation using an Illumina Nextera XT DNA Library Preparation Kit prior to 2×100bp paired-end sequencing via Illumina HiSeq 4000 technology. Raw sequence reads were quality controlled with BBTools (Bushnell 2014) and assembled with MegaHit v. 1.02 (Li *et al.*, 2016). The OHK43 genome bin was recovered via differential coverage binning with MetaBAT (Kang *et al.*, 2015). Completeness and contamination/redundancy were determined with CheckM v1.1.2 (Parks *et al.*, 2015). The genome was uploaded to RAST v2.0 for annotation and characterization (Aziz *et al.*, 2008). Presence or absence of metabolic pathways of interest was predicted using MetaPOAP v1.0 (Ward *et al.*, 2018). Taxonomic assignment was determined with GTDB-Tk v1.2 (Parks *et al.*, 2018, 2020; Chaumeil *et al.*, 2019).

### Organismal Phylogenies

Concatenated ribosomal phylogenies were constructed following methods from Hug *et al.* (2016). Members of the *Thermosynechococcaceae* and outgroups were identified using GTDB (Chaumeil *et al.*, 2019) and their genomes downloaded from the NCBI WGS and Genbank databases. Ribosomal protein sequences were extracted using the tblastn function of BLAST+ (Camacho *et al.*, 2009) and aligned with MUSCLE (Edgar, 2004). Trees were built with RAxML v.8.2.12 (Stamatakis, 2014) on the Cipres science gateway (Miller *et al.*, 2010). Transfer bootstrap support values were determined with BOOSTER (Lemoine *et al.*, 2018). Visualization of trees was performed with the Interactive Tree of Life Viewer (Letunic and Bork, 2016).

### Genome comparison

We compared the core- and pangenomes of *Thermosynechococcus* at the genus level, family level and with a sub-sample of organisms across the GTDB defined class Cyanobacteriia. ProteinOrtho (Lechner *et al.*, 2011) was used for the identification of Conserved Likely Orthologous Groups (CLOG) and analysis.

At the genus level, ten *Thermosynechococcus* strains from varying hot spring environments (Table 1 and 2) were compared. The genomic data from ten currently available sequences of *Thermosynechococcus* strains *T.* sp. CL1/1-2178 (CL1), *T. elongatus* BP1/1-2178 (*elongatus* BP1)*, T*. *vulcanus* NIES2134/1-2178 (*vulcanus*)*, T.* sp. NK55a/1-2022 (NK55a), *T. elongatus* PKUAC-SCTE542 (*elongatus* PKUAC), *T.* sp. M55_K2018_012 (Tattapani 1), *T.* sp. M98_K2018_005 (Tattapani 2), *Ca. T.* sp. J003 (J003), *T.* sp. M3746_W2019_013 (Jinata 2) and *Ca. T.* sp. OHK43 (OHK43) were compared using ProteinOrtho (Lechner *et al.*, 2011), BLAST (Altschul *et al.*, 1990), and FeGenie (Garber *et al.*, 2019). Due to low completeness of the genome, we excluded the eleventh available species (*T.* sp. M46_R2017_013) from this analysis. Phylogenetic relationships between these strains were established using concatenated ribosomal protein phylogenies following Hug *et al.* (2016), taxonomic classifications with GTDB-tk (Chaumeil *et al.*, 2019) and average nucleotide identities (Rodriguez-R and Konstantinidis, 2017).

**Table 2:** Genus level average nucleotide identities (ANI) and genome sizes (diagonal). Additionally, genomes sizes for family level species are added in the lower left.

| Genome Size | Ca. *T. nakabusesensis* Jinata | Ca. *T. nakabusesensis* OHK | *T. elongatus* BP-1 | *T. elongatus* PKUAC-SCTE542 | *T. sp* CL1 | *T. nakabusesensis* NK55a | *T. vulcanus* NIES2134 | *T. sp.* M3746_W2019_013 | *T. sp.* M46_R2017_013 | *T. sp.* M55_K2018_012 | *T. sp.* M98_K2018_005 |
| --- | --- | --- | --- | --- | --- | --- | --- | --- | --- | --- | --- |
| Ca. *T. nakabusesensis* Jinata | **2.31 MB** | 99.70 | 92.65 | 86.82 | 87.53 | 99.68 | 92.59 | 99.76 | 83.92 | 87.85 | 87.85 |
| Ca. *T. nakabusesensis* OHK |  | **2.25 MB** | 92.58 | 86.85 | 87.52 | 99.84 | 92.55 | 99.76 | 83.95 | 87.88 | 87.87 |
| *T. elongatus* BP-1 |  |  | **2.59 MB** | 86.65 | 87.34 | 92.53 | 99.14 | 92.56 | 83.86 | 87.63 | 87.61 |
| *T. elongatus* PKUAC-SCTE542 |  |  |  | **2.64 MB** | 89.95 | 86.81 | 86.63 | 86.88 | 85.45 | 90.54 | 90.58 |
| *T. sp* CL1 |  |  |  |  | **2.64 MB** | 87.50 | 87.32 | 87.55 | 85.51 | 92.27 | 92.26 |
| *T. nakabusesensis* NK55a | Genome Sizes for GTDB representative family level species:<br>*Acaryochloris marina* MBIC11017 8.36 MB<br>*Acaryochloris* sp. CCMEE 5410 7.87 MB<br>*Cyanothece* sp. PCC 7425 5.78 MB<br>*Synechococcus lividus* PCC 6715 2.65 MB<br>*Synechococcus* sp. PCC 6312 3.72 MB<br>*Acaryochloris* RCC 1774 5.93 MB |  |  |  |  | **2.51 MB** | 92.45 | 99.69 | 83.92 | 87.84 | 87.82 |
| *T. vulcanus* NIES2134 |  |  |  |  |  |  | **2.57 MB** | 92.51 | 83.79 | 87.55 | 87.56 |
| *T. sp.* M3746_W2019_013 |  |  |  |  |  |  |  | **2.38 MB** | 83.99 | 87.87 | 87.90 |
| *T. sp.* M46_R2017_013 |  |  |  |  |  |  |  |  | **2.39 MB** | 86.03 | 86.06 |
| *T. sp.* M55_K2018_012 |  |  |  |  |  |  |  |  |  | **2.39 MB** | 99.92 |
| *T. sp.* M98_K2018_005 |  |  |  |  |  |  |  |  |  |  | **2.37 MB** |

For the analysis at the family level, we included 6 more species that appear as representative strains at the *Thermosynochoccaceae* family level in the GTDB. The representative GTDB strains include *Acaryochloris marina* MBIC11017, *Cyanothece* sp. PCC 7425, *Acaryochloris* sp. CCMEE 5410, *Synechococcus* sp. PCC 6312, *Synechococcus lividus* PCC 6715 and *Acaryochloris* sp. RCC1774. This resulted in a total of 16 strains for family level analysis.

At the class level we used the ten genus level strains and additionally 16 species, 15 after (Beck *et al.*, 2012) and in addition one *Gloeobacter* species (*G. kilaueensis* JS-1). The class level analysis includes all of the genus level strains, and some family and class level strains. Here the goal was to test the coherence of our analysis parameters when compared to previous studies of cyanobacterial core- and pangenomes (Beck *et al.*, 2012, 2018). We acknowledge that the distribution of species within the families of the class is not even, however this comparison was used to verify our methods and the stability of the class level core. This resulted in a total of 26 genomes for the class level analysis.

To compare the appearance of genes related to iron uptake and regulation we used FeGenie (Garber *et al.*, 2019) with standard parameters. We used results from ProteinOrtho (Lechner *et al.*, 2011) for further analysis of the core, shared, unique, TS (Thermosynechococcus) core and TS shared clusters. Proteinortho was applied such that the output included also singleton clusters (only containing a single protein) and with an algebraic connectivity of 0 as a measure for the structure of the orthologous clusters. We did not obtain many differences when running ProteinOrtho on our data using a value of 0 or 0.1 (default) for the algebraic connectivity, however, a value of 0 resulted in slightly larger clusters for the core and a few less singletons. A comparison of results from FeGenie and ProteinOrtho resulted in similar gene clusters for iron related genes such that our results here are based on both program outputs.

## Results and Discussion

### Phylogeny of the Thermosynechococcus and the species proposal T. nakabusensis

The *Thermosynechococcus* genus is phylogenetically coherent within the Cyanobacteria (Figure 1) and the genome sizes of genus members are similar to one another (Table 2). Based on similarity observed with ANI and GTDB-tk (Table 2), 4 species are present within the *Thermosynechococcus* genus: *T. elongatus* BP1 and *T. vulcanus* belonging to one species, *T.* NK55a, *Ca. T.* sp. J003, *T.* sp. M3746_W2019_013 (Jinata 2) and *Ca. T.* sp. OHK43 (OHK43) genomes belonging to a second species, Tattapani 1 and Tattapani 2 belonging to a third species, and *T.* CL1 and *T. elongatus* PKUAC-SCTE542 as one species each. For *T.* sp. M46_R2017_013 (Shivlinga) the species designation remains unresolved due to low completeness. For the species including *T.* NK55a, J003 and Jinata 2, and OHK43 genomes we propose the name *Thermosynechococcus nakabusensis* after the first and so far only isolated organism which originates from Nakabusa hot spring in Nagano Prefecture, Japan.

**Figure 1:**
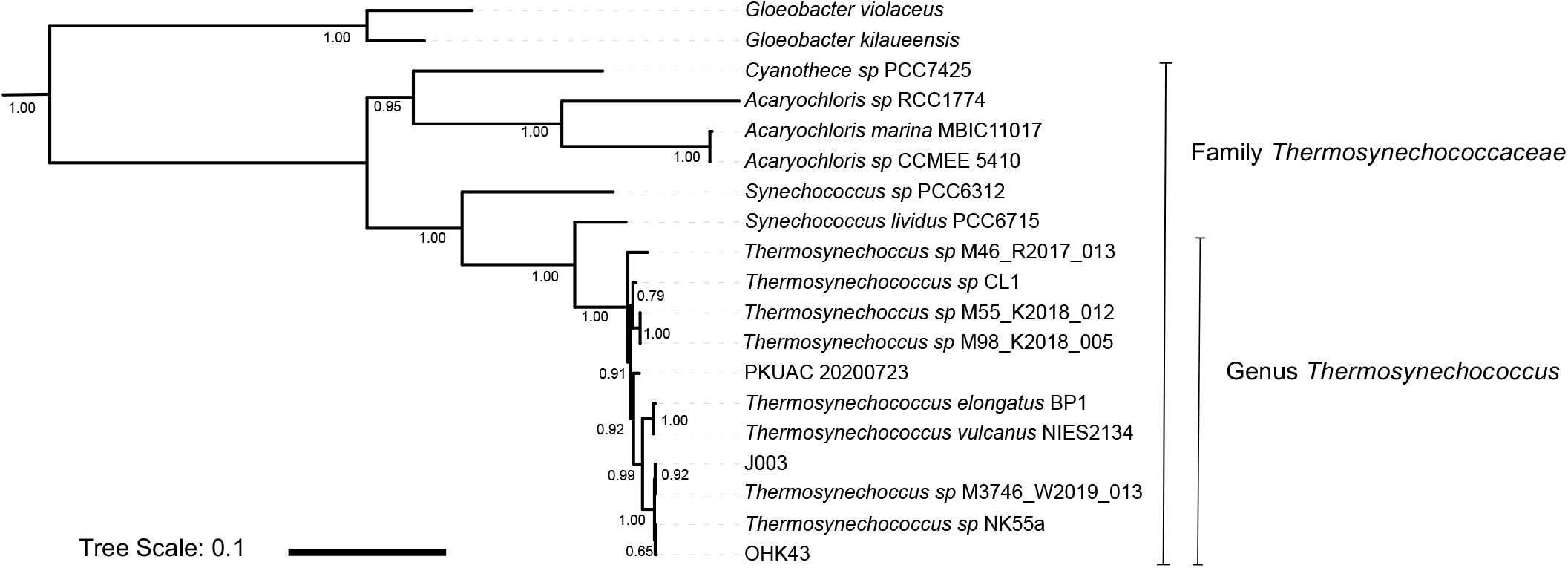
*Thermosynechococcaceae* phylogeny built with concatenated ribosomal proteins. Branch supports are derived from bootstrapping with BOOSTER and the tree scale bar indicates substitutions per nucleotide site.

### Genus and family level comparison of the genus *Thermosynechococcus* and the family *Thermosynochococcaceae*

Comparing conserved likely orthologous groups (CLOGs), we analyzed i) the core-genomes: those CLOGs shared by all genomes in an analysis, ii) the shared CLOGs: those shared by at least 2 but not all of the genomes in the analysis, and iii) unique CLOGs: those CLOGs that are unique to a single genome (Table 3). The *Thermosynechococcus* genus specific core (**core TS**) comprises CLOGs shared by all ten genus level genomes that are not present in any other species, and the *Thermosynechococcus* genus-specific shared CLOGs (**shared TS**) corresponds to CLOGs that are shared by at least two and at most nine genus level genomes.

**Table 3 -.** Numbers of CLOGs per grouping and phylogenetic level. *note that the number of genus specific core CLOGs increases due to the exclusion of some family level genomes at the class level.

|  | Genus (10 strains) | Family (16 strains) | Class (26 strains) |
| --- | --- | --- | --- |
| Core of all genomes analyzed | 1737 | 1225 | 723 |
| *Thermosynechococcus* genus specific core | - | 14 | 67 |
| Uniquely shared between 7 *Thermosynechococcus* (not including J003/Jinata2/OHK43) | 1 | 0 | 0 |
| Uniquely shared between J003, Jinata 2 and OHK43 only (high iron organisms) | 1 | 1 | 1 |

Comparing the genomes of the ten genus members, the protein core is made up of 1737 CLOGs and contains 66 to 75 % of the putative protein coding genes in a genome. This value is different from a recent analysis (Cheng *et al.*, 2020), who compared 5 genomes available at that time and found a core of 1264 CLOGs. The higher value observed here is attributed to the analysis of a revised *T. elongatus* PKUAC-SCTE542 genome which had a much lower quantity of pseudogenes in comparison to the previously available version.

The percentage of the *Thermosynechococcus* genomes which belongs to the genus core is higher than in other comparisons of organisms analyzed at this taxonomic level, for example Wu *et al.* (2018) found that the core genes of species belonging to the genus *Comamonas* account for 18 – 33 % of all genes, and Barajas *et al.* (2019) reported that the core genome within the genus *Streptococcus* ranges in size from 9.6 % to 24 %. At the species level, Reno *et al.* (2009) found that 69 – 79 % of genes made up the core in *S. islandicus* strains. The high proportion of CLOGs which make up the core leaves few unique CLOGs for each genome and between genus members, and only slight variations in genome content between genomes are observed: focusing on the organisms from high ferrous iron springs compared to low iron springs for example, only one CLOG is specific to the seven genomes that do not include J003, Jinata 2 and OHK43 and 1 CLOG is specific to J003, Jinata 2 and OHK43 (Table 3, supplementary Figure 1).

Golicz *et al.* (2020) suggested that the size of a pangenome is related to organismal lifestyle, with sympatric organisms having open pangenomes with many accessory genes and allopatric organisms having more closed and conserved pangenomes. The *Thermosynechococcus* genus could thus be considered as allopatric as they have a comparatively large core and few shared and unique genes. Since all *Thermosynechococcus* in this study are found in hot springs – which typically have reduced microbial diversity in comparison to other environments such as soils (Ward *et al.*, 1998) – the conserved core and small pangenome of the *Thermosynechococcus* genus may reflect a more limited opportunity for lateral gene transfer, which in tuxrn could lead to less opportunity for lateral gene transfer and smaller genomes.

Chen *et al.* (2020) also noted that differences in horizontal gene transfer (HGT) are related to genome size with smaller genomes showing less HGT and larger genomes having a greater probability that HGT occurred. They also found that hot spring cyanobacteria specifically have smaller genome sizes and less HGT into the genome. Excluding massive gene loss within the *Thermosynechoccus* genus, it is tempting to speculate that the proportionally large *Thermosynechoccus* genus core might be indicative of a more ancient gene repertoire in hot spring cyanobacteria, with other cyanobacteria gaining more functionality through HGT over evolutionary timescales. Although still tentative, the observation that thermotolerance is phylogenetically scattered across the cyanobacterial tree of life and occurs mostly in organisms comprising smaller genomes is in line with this hypothesis of substantial gene gain by HGT in many cyanobacterial lineages and a small ancestral core.

In contrast to the genus level where genome size varies from 2.25 MB to 2.64 MB, at the family level it varies up to 8.36 MB (*Acaryocholoris marina* MBIC11017, Table 2). Running Proteinortho analyses with the ten genus level *Thermosynechococcus* and the six family level sequences, the shared genes, not core genes, comprise a larger percentage of the genomes for smaller genomes, while unique genes are abundant in larger genomes (Figure 2a). The overall core is reduced by about 30%, from 1737 to 1225 CLOGs.

**Figure 2:**
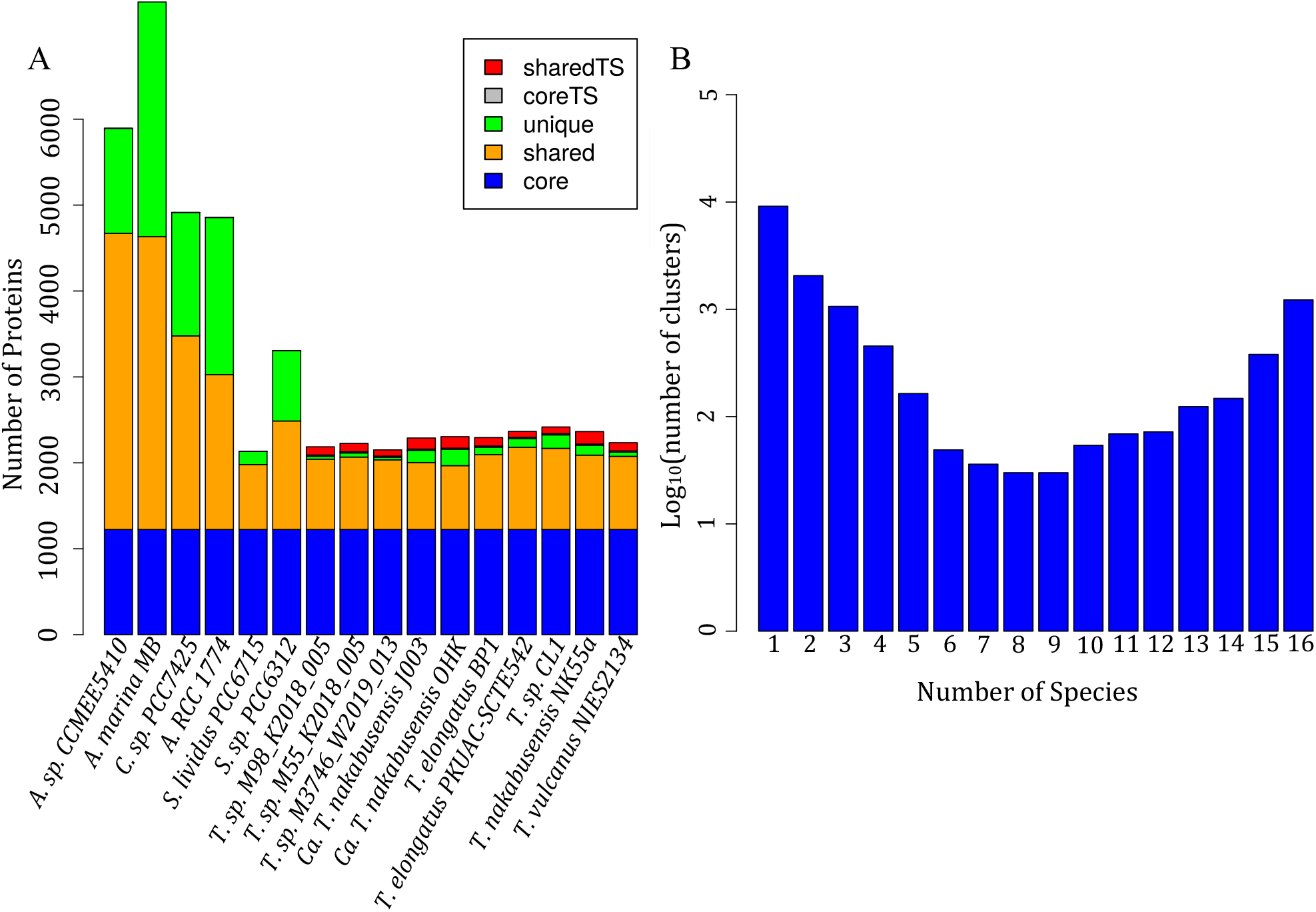
(A) Thermosynechococcaceae family level comparison of core- and pangenomes, (B) number of CLOGs observed at the family level in relationship to the number of genomes in the analysis. coreTS indicates CLOGs found in all genus level genomes, sharedTS are CLOGs found in at least 2 but not all genus level genomes and no other genomes. Full species names as mentioned above.

At the family level, the number of CLOGs shared between the seven *Thermosynechococcus* genus members that do not include J003, Jinata 2 and OHK43 (the strains from higher iron environments) is reduced from one (found when analyzing at the genus level) to zero, showing that this CLOG is found in other closely related members at the family level. However, the number of CLOGs found only in J003 and OHK43 changes from 23 to 20 CLOGs, highlighting that these are unique even at the family level (SI Table 2).

### Genomic differentiation of the *Thermosynechococcus* genus from other cyanobacteria

At the class level the number of *Thermosynechococcus* genus level core CLOGs increases when compared to the family level, and this is due to the exclusion of some family level species at this level (in our analysis the class level comparison was made specifically in reference to a previous study (Beck *et al.* 2012) for comparative purposes; see Methods and Table 3). Consistent with this former study, the overall core decreases by almost 60% when class level representatives are added (from 1737 core CLOGs at the genus level to 723 core CLOGs at the class level). This is expected and in line with the larger analyses of microbial genomes which have shown that the continued addition of taxonomic diversity in an analysis leads to increasingly smaller cores (Charlebois and Doolittle, 2004; Lapierre and Gogarten, 2009). Moreover, Beck *et al.* (2018) suggested that the clustering of CLOGs depends on the data analyzed including variability in genome size as well as phylogenetic distance between the analyzed genomes. Overall these results conform with Beck *et al.* (2012), with slight variations attributed to the different genomes used and the different orthology methods and parameters used for each analysis. Finally, we confirm that the class level core is stable when adding a higher diversity of organisms - similar to Beck *et al.* (2018) - as the number of core family CLOGs only slightly decreases when adding more species at the class level (660 core CLOGs were obtained with 16 strains in Beck *et al.* (2012) and 621 core CLOGs with 77 strains in Beck *et al.* (2018)).

Investigating the genes which may differentiate the *Thermosynechococcus* genus from other cyanobacteria, we found that 67 CLOGs appear in all *Thermosynechococcus* but not in any other cyanobacteria. 37 of these are hypothetical proteins of unknown functions and could be of interest to further biochemical studies to understand the adaptations of the genus. These genes may code for proteins that make the *Thermosynechococcus* unique from other cyanobacteria and appear to be related to transcriptional regulation, transporters and membrane proteins (e.g. acetyltransferases, glycosidases and ATPases; Supplementary Table 1). Since all the *Thermosynechococcus* analyzed are from hot springs, these genes potentially provide some basis for that lifestyle.

### Adaptation of Thermosynechococcus to their respective environments

It is important to note that although the spring source water properties differ, the source water is not necessarily where the DNA or the isolated organism originated. Additionally, hot spring flow paths frequently change course, so even if an organism is isolated at one time from one location, it may have been previously growing under a different set of conditions, leading to confusion between the relationship between genotype and phenotype. Acknowledging this, the gross qualities of the hot springs analyzed here differ significantly in pH and the type of reductant present (*e.g.* iron vs. sulfide) (Table 1) implying that organisms inhabiting them experience different environments. We were especially interested in those CLOGs that are shared between the strains from environments with elevated iron concentrations (the genomes J003, Jinata 2 and OHK43) but which are not present in any other cyanobacteria. Previous studies have shown that some cyanobacteria express higher levels of genes involved in iron ion homeostasis in iron limiting conditions (Cheng and He, 2014), and we investigated the presence or absence of iron related gene products in *Thermosynechococcus* compared to other cyanobacteria using FeGenie and BLAST comparisons. Only one CLOG is uniquely shared between J003, Jinata 2 and OHK43 amongst the genus members. Analyzed at the class level, 19 CLOGs are uniquely shared between Jinata 1 and OHK43. It is notable that this number is higher than the CLOGs shared between all three strains, and that these 19 CLOGs do not appear in the second strain from Jinata (Jinata 2) or in the fourth strain of the same species (NK55a). Two CLOGs are uniquely shared between OHK43 and Jinata 2, and six CLOGs are uniquely shared between J003 and Jinata 2. Of these genes, some show high partial identity but low coverage matches with genes from other *Thermosynechococcus,* especially *T.* sp. NK55a (supplementary Table 2). With our current understandings after considering results from BLAST and FeGenie (SI Table 2) none of the CLOGs shared by the organisms from high iron hot springs comprises genes that could explain adaptation to elevated iron concentrations. From these analyses we conclude that there is no sequence resolvable genomic signature specific to strains from Jinata and OHK hot springs related to iron tolerance or oxidative stress response.

*Thermosynechococcus* lack genes coding for ferrous iron transport and uptake proteins EfeB, EfeO and EfeU, the metal transport gene ZupT, the cellular iron storage protein Bfr, and the iron regulator active under iron limiting conditions PfsR (Table 4). In all cases the same genes encoding proteins related to ferrous iron uptake (FeoA, FeoB, YfeA and YfeB), ferric iron uptake or transport (ExbD, FutA, FutB and FutC), siderophore iron acquisition (FpvD), metal ion binding (Ho1 and Ho2) and iron starvation acclimation (IsiA) are present (Table 4).

**Table 4:** Genes known to be involved in iron regulation within the class Cyanobacteriia and their presence in the *Thermosynechococcus* genus. Presence or absence of genes was confirmed with BLAST searches and FeGenie.

| Protein | Pfam identifier | Function | Present in *Thermosynechococcus* | Reference |
| --- | --- | --- | --- | --- |
| FeoA | PF04023 | ferrous iron uptake | Yes, all | Lau *et al.*, 2016 |
| FeoB | PF07664 | ferrous iron uptake | Yes, all | Lau *et al.*, 2016 |
| YfeA | PF01297 | ferrous iron uptake | Yes, all | Toulza *et al.*, 2012 |
| YfeB | PF00005 | ferrous iron uptake | Yes, all | Toulza *et al.*, 2012 |
| FpvD | PF00005 | siderophore iron acquisition | Yes, all | Brillet *et al.*, 2012 |
| ExbD | PF02472 | ferric iron uptake | Yes, all | Jiang *et al.*, 2015 |
| PfsR | PF00440 | iron regulator under iron limiting conditions | No, but other cyanobacteria | Cheng and He, 2014 |
| FutA | PF13416 | ABC-type ferric iron transport | Yes, all | Katoh *et al.*, 2001 |
| FutB | PF00528 | ABC-type ferric iron transport | Yes, all | Katoh *et al.*, 2001 |
| FutC | PF00005 | ABC-type ferric iron transport | Yes, all | Katoh *et al.*, 2001 |
| EfeB | PF04261 | ferrous iron transport | No, but other cyanobacteria | Lau *et al.*, 2016 |
| EfeO | PF13473 | ferrous iron transport | No, but other cyanobacteria | Lau *et al.*, 2016 |
| EfeU | PF03239 | ferrous iron uptake | No, but other cyanobacteria | Lau *et al.*, 2016 |
| ZupT | PF02535 | metal transport (including ferrous iron) | No, but other cyanobacteria | Lau *et al.*, 2016 |
| Bfr | PF00210 | cellular iron storage | No, but other cyanobacteria | Keren *et al.*, 2004; Cheng and He, 2014 |
| IsiA | PF00421 | iron starvation acclimation | Yes, all | Cheng and He, 2014 |
| Ho1 | PF01126 | metal ion binding | Yes, all | Cheng and He, 2014 |
| Ho2 | PF01126 | metal ion binding | Yes, all | Cheng and He, 2014 |

In the absence of a resolvable genetic signature for iron tolerance, we recall that Ionescu *et al.* (2014) proposed that by simply increasing photosynthetic rate and oxygen production, cyanobacteria might protect themselves from ferrous iron by promoting its precipitation at some distance from the cell (Ionescu *et al.*, 2014). In line with this observation, is worthwhile to note that the biomass accumulation in high iron environments like Jinata hot spring is appreciable, with co-occurring visible biomass and dissolved oxygen concentrations that are elevated above what is expected from atmospheric solubility, indicating active water oxidizing photosynthesis (Ward *et al.*, 2019).

### The conserved genomic core of *Thermosynechococcus* in relationship to environmental distribution is unique

In addition to uncovering a highly conserved genome core in a group of organisms with significant environmental distribution, our work is also relevant to historical proliferation of cyanobacteria, since some modern-day hot springs and their biogeochemistries can be used as historical process analogues (Brown *et al.*, 2005, 2007; Ward *et al.*, 2019). Considering contemporary environments, the analysis of *Thermosynechococcus* also provides insight into island biogeography of microbes. Ionescu *et al.* (2010) observed that the speciation patterns of microorganisms are shaped by local community structures and environmental influences, and Bahl *et al.* (2011) additionally suggest a positive correlation between geographic and genetic distance. Papke *et al.* (2003) found that isolated environments such as geothermal springs may lead to evolutionary divergence of closely related *Thermosynechococcus* strains due to island effects, similar to analyses by Whitaker *et al.* (2003), who found similar trends of divergence in hypterthermophilic archaea. Our analysis suggests that geographically widespread organisms belonging to the genus inhabit hot springs with varying geochemistries without genomically recognizable adaptations specific to their site of origin. Instead, the finding of highly conserved genomes within the genus, and furthermore, that the genetic content of the genus is not markedly different from other cyanobacteria, implies that the genus is inherently flexible and able to grow in the geochemical regimes studied. The large portion of shared genes within the genus provides a genetic basis for the lack of correlation between geographic and genetic distances within the genus found by Papke *et al.* (2003). Apparently, *Thermosynechococcus* is environmentally promiscuous, and have fewer restrictive requirements concerning their distribution. This is in contrast to other groups of organisms, for example the analyzed by Reno *et. al.* (2009) who found that the obligately thermoacidophilic *S. islandicus* archaea show a core and pangenome shaped by their geographical distribution.

## Outlook

Based on the genome comparisons presented here, a viability test of isolates in environments other than those of their origin is suggested as future work. For example, genus level iron tolerance experiments are proposed to test if strains from low-iron environments can withstand elevated iron. In a similar way, thermotolerance of these organisms could also be investigated. This could help us understand if *Thermosynechococcus* is indeed less restrictive with respect to their geochemical requirements or to identify mechanisms unresolvable by the CLOG approach that account for the geographical distribution.

## Data availability for newly described strains

The Whole Genome Shotgun project for OHK43 has been deposited at DDBJ/ENA/GenBank under the accession JACOMP000000000. The version described in this paper is version JACOMP010000000.

## Supplemental Material

SI Figure 1 – class and genus level comparison of core- and pan-genomes; distribution of number of CLOGs with number of genomes at class and genus level. Core TS corresponds to the CLOGs found in all *Thermosynechococcus* genus members, and shared TS corresponds to CLOGs that are shared by at least 2 and at most 6 genus level genomes.

SI Table 1 – *Thermosynechococcus* genus core of CLOGs obtained from analysis at class level and their corresponding BLAST hit annotations when using the *T. vulcanus* protein sequences as query. The annotation is derived from the top hit (either the *T. vulcanus* annotation, or the multispecies annotation). The highlighted CLOGs are unique both at family and at class level comparisons.

SI Table 2 – Jinata and OHK specific CLOGs at class level with coverage and identity scores

## Acknowledgements

LMW was supported by a Simons Foundation Postdoctoral Fellowship in Marine Microbial Ecology. Metagenomic sequencing of OHK was supported by NSF grant #OISE 1639454. SEM acknowledges support from the Astrobiology Center Program of the National Institutes of Natural Sciences (grant no. AB311013). We would also like to thank the group around M. Daroch for providing us with an updated version for the *T. elongatus* PKUAC-SCTE542 genome which increased the strain’s coherence compared to previous versions of the genome.

## Notes

### Competing Interest Statement

The authors have declared no competing interest.

### Summary of Updates

We have added the doi of the final accepted version after the abstract. This version also includes additional data.

## References

Alcorta, J., Alarcón-Schumacher, T., Salgado, O., and Díez, B. (2020) Taxonomic Novelty and Distinctive Genomic Features of Hot Spring Cyanobacteria. Front Genet 11:.

Allen, J.F., Thake, B., and Martin, W.F. (2019) Nitrogenase Inhibition Limited Oxygenation of Earth’s Proterozoic Atmosphere. Trends in Plant Science 24: 1022–1031.

Altschul, S.F., Gish, W., Miller, W., Myers, E.W., and Lipman, D.J. (1990) Basic local alignment search tool. Journal of Molecular Biology 215: 403–410.

Aziz, R.K., Bartels, D., Best, A., DeJongh, M., Disz, T., Edwards, R.A., et al. (2008) The RAST Server: Rapid annotations using subsystems technology. BMC Genomics 9: 75.

Bahl, J., Lau, M.C.Y., Smith, G.J.D., Vijaykrishna, D., Cary, S.C., Lacap, D.C., et al. (2011) Ancient origins determine global biogeography of hot and cold desert cyanobacteria. Nature Communications 2: 1–6.

Barajas, H.R., Romero, M.F., Martínez-Sánchez, S., and Alcaraz, L.D. (2019) Global genomic similarity and core genome sequence diversity of the Streptococcus genus as a toolkit to identify closely related bacterial species in complex environments. PeerJ 2019: e6233.

Beck, C., Knoop, H., Axmann, I.M., and Steuer, R. (2012) The diversity of cyanobacterial metabolism: genome analysis of multiple phototrophic microorganisms. BMC genomics 13: 56.

Beck, C., Knoop, H., and Steuer, R. (2018) Modules of co-occurrence in the cyanobacterial pan-genome reveal functional associations between groups of ortholog genes. PLOS Genetics 14: e1007239.

Brillet, K., Ruffenach, F., Adams, H., Journet, L., Gasser, V., Hoegy, F., et al. (2012) An ABC Transporter with Two Periplasmic Binding Proteins Involved in Iron Acquisition in Pseudomonas aeruginosa. ACS Chemical Biology 7: 2036–2045.

Callieri, C., Slabakova, V., Dzhembekova, N., Slabakova, N., Peneva, E., Cabello-Yeves, P.J., et al. (2019) The mesopelagic anoxic Black Sea as an unexpected habitat for Synechococcus challenges our understanding of global “deep red fluorescence.” ISME Journal 13: 1676–1687.

Camacho, C., Coulouris, G., Avagyan, V., Ma, N., Papadopoulos, J., Bealer, K., and Madden, T.L. (2009) BLAST+: Architecture and applications. BMC Bioinformatics 10: 1–9.

Canfield, D.E. (1998) A new model for Proterozoic ocean chemistry. Nature 396: 450.

Charlebois, R.L. and Doolittle, W.F. (2004) Computing prokaryotic gene ubiquity: Rescuing the core from extinction. Genome Research 14: 2469–2477.

Chaumeil, P.-A., Mussig, A.J., Hugenholtz, P., and Parks, D.H. (2019) GTDB-Tk: a toolkit to classify genomes with the Genome Taxonomy Database. Bioinformatics 36: 1925–1927.

Chen, M.Y., Teng, W.K., Zhao, L., Hu, C.X., Zhou, Y.K., Han, B.P., et al. (2020) Comparative genomics reveals insights into cyanobacterial evolution and habitat adaptation. ISME J 1–17.

Cheng, D. and He, Q. (2014) PfsR Is a Key Regulator of Iron Homeostasis in Synechocystis PCC 6803. PLoS ONE 9: e101743.

Cheng, Y.-I., Chou, L., Chiu, Y.-F., Hsueh, H.-T., Kuo, C.-H., and Chu, H.-A. (2020) Comparative Genomic Analysis of a Novel Strain of Taiwan Hot-Spring Cyanobacterium Thermosynechococcus sp. CL-1. Frontiers in Microbiology 11: 82.

Edgar, R.C. (2004) MUSCLE: a multiple sequence alignment method with reduced time and space complexity. BMC Bioinformatics 5:113

Fennel, K., Follows, M., and Falkowski, P.G. (2005) The co-evolution of the nitrogen, carbon and oxygen cycles in the proterozoic ocean. American Journal of Science 305: 526–545.

Fischer, W.W., Hemp, J., and Valentine, J.S. (2016) How did life survive Earth’s great oxygenation? Current Opinion in Chemical Biology 31: 166–178.

Garber, A.I., Nealson, K.H., Okamoto, A., McAllister, S.M., Chan, C.S., Barco, R.A., and Merino, N. (2019) FeGenie: a comprehensive tool for the identification of iron genes and iron gene neighborhoods in genomes and metagenome assemblies. bioRxiv 777656.

Golicz, A.A., Bayer, P.E., Bhalla, P.L., Batley, J., and Edwards, D. (2020) Pangenomics Comes of Age: From Bacteria to Plant and Animal Applications. Trends in Genetics 36: 132–145.

Guo, Q., Pang, Z., Wang, Y., and Tian, J. (2017) Fluid geochemistry and geothermometry applications of the Kangding high-temperature geothermal system in eastern Himalayas. Applied Geochemistry 81: 63–75.

Hao, J., Knoll, A.H., Hazen, R.M., Daniel, I., Huang, F., and Schieber, J. (2020) Cycling Phosphorus on the Archean Earth: Part II. Phosphorus Limitation on Primary Production in Archean Ecosystems Pattern-forming abiotic reactions during organic diagenesis View project Deep Time Data Infrastructure View project Cycling phosphorus on the Archean Earth: Part II. Phosphorus limitation on primary production in Archean ecosystems ScienceDirect.

Hsueh, H.T., Chu, H., and Chang, C.C. (2007) Identification and Characteristics of a Cyanobacterium Isolated from a Hot Spring with Dissolved Inorganic Carbon. Environmental Science & Technology 41: 1909–1914.

Hug, L.A., Baker, B.J., Anantharaman, K., Brown, C.T., Probst, A.J., Castelle, C.J., et al. (2016) A new view of the tree of life. Nature Microbiology 1: 1–6.

Ichikuni, M. and Kikuchi, K. (1972) Retention of boron by travertines. Chemical Geology 9: 13–21.

Ionescu, D., Hindiyeh, M., Malkawi, H., and Oren, A. (2010) Biogeography of thermophilic cyanobacteria: insights from the Zerka Ma’in hot springs (Jordan). FEMS Microbiol Ecol 72: 103–113.

Ionescu, D., Buchmann, B., Heim, C., Häusler, S., de Beer, D., and Polerecky, L. (2014) Oxygenic photosynthesis as a protection mechanism for cyanobacteria against iron-encrustation in environments with high Fe2+ concentrations. Frontiers in Microbiology 5: 459.

Jiang, H.-B., Lou, W.-J., Ke, W.-T., Song, W.-Y., Price, N.M., and Qiu, B.-S. (2015) New insights into iron acquisition by cyanobacteria: an essential role for ExbB-ExbD complex in inorganic iron uptake. The ISME journal 9: 297.

Kang, D.D., Froula, J., Egan, R., and Wang, Z. (2015) MetaBAT, an efficient tool for accurately reconstructing single genomes from complex microbial communities. PeerJ 2015: e1165.

Katoh, H., Hagino, N., and Ogawa, T. (2001) Iron-Binding Activity of FutA1 Subunit of an ABC-type Iron Transporter in the Cyanobacterium Synechocystis sp. Strain PCC 6803, Oxford Academic.

Kaushal, G., Kumar, J., Sangwan, R.S., and Singh, S.P. (2018) Metagenomic analysis of geothermal water reservoir sites exploring carbohydrate-related thermozymes. Int J Biol Macromol 119: 882–895.

Keren, N., Aurora, R., and Pakrasi, H.B. (2004) Critical roles of bacterioferritins in iron storage and proliferation of cyanobacteria. Plant Physiology 135: 1666–1673.

Knoll, A.H. (2006) Cyanobacteria and Earth History Diverse aspects of testate amoebae evolution View project The Co-Evolution of the Geo-and Biospheres: An Integrated Program for Data-Driven Abductive Discovery in Earth Sciences View project.

Lapierre, P. and Gogarten, J.P. (2009) Estimating the size of the bacterial pan-genome. Trends in Genetics 25: 107–110.

Lau, C.K.Y., Krewulak, K.D., and Vogel, H.J. (2016) Bacterial ferrous iron transport: the Feo system. FEMS microbiology reviews 40: 273–298.

Lechner, M., Findeiß, S., Steiner, L., Marz, M., Stadler, P.F., and Prohaska, S.J. (2011) Proteinortho: detection of (co-) orthologs in large-scale analysis. BMC bioinformatics 12: 124.

Lemoine, F., Entfellner, J.-B.D., Wilkinson, E., Correia, D., Felipe, M.D., De Oliveira, T., and Gascuel, O. (2018) Renewing Felsenstein’s phylogenetic bootstrap in the era of big data. Nature 556: 452–456.

Letunic, I. and Bork, P. (2016) Interactive tree of life (iTOL) v3: an online tool for the display and annotation of phylogenetic and other trees. Nucleic Acids Res 44: W242–W245.

Li, D., Luo, R., Liu, C.M., Leung, C.M., Ting, H.F., Sadakane, K., et al. (2016) MEGAHIT v1.0: A fast and scalable metagenome assembler driven by advanced methodologies and community practices. Methods 102: 3–11.

Lyons, T.W., Reinhard, C.T., and Planavsky, N.J. (2014) The rise of oxygen in Earth’s early ocean and atmosphere. Nature 506: 307.

Maity, J.P., Liu, C.-C., Nath, B., Bundschuh, J., Kar, S., Jean, J.-S., et al. (2011) Biogeochemical characteristics of Kuan-Tzu-Ling, Chung-Lun and Bao-Lai hot springs in southern Taiwan. Journal of Environmental Science and Health, Part A 46: 1207–1217.

Mei-hua, W., Junhao, W., and Tingshan, T. (2015) Study of the Scaling Trend of Thermal Groundwater in Kangding County of Sichuan Province. World Geothermal Congress 2015 11.

Miller, M.A., Pfeiffer, W., and Schwartz, T. (2010) Creating the CIPRES Science Gateway for inference of large phylogenetic trees. In 2010 gateway computing environments workshop (GCE). Ieee, pp. 1–8.

Nakagawa, T. and Fukui, M. (2002) Phylogenetic characterization of microbial mats and streamers from a Japanese alkaline hot spring with a thermal gradient. The Journal of General and Applied Microbiology 48: 211–222.

Nakamura, Y., Kaneko, T., Sato, S., Ikeuchi, M., Katoh, H., Sasamoto, S., et al. (2002) Complete genome structure of the thermophilic cyanobacterium Thermosynechococcus elongatus BP-1. DNA research 9: 123–130.

Papke, R.T., Ramsing, N.B., Bateson, M.M., and Ward, D.M. (2003) Geographical isolation in hot spring cyanobacteria. Environmental Microbiology 5: 650–659.

Parks, D.H., Chuvochina, M., Chaumeil, P.A., Rinke, C., Mussig, A.J., and Hugenholtz, P. (2020) A complete domain-to-species taxonomy for Bacteria and Archaea. Nature Biotechnology 1–8.

Parks, D.H., Chuvochina, M., Waite, D.W., Rinke, C., Skarshewski, A., Chaumeil, P.A., and Hugenholtz, P. (2018) A standardized bacterial taxonomy based on genome phylogeny substantially revises the tree of life. Nature Biotechnology 36: 996.

Parks, D.H., Imelfort, M., Skennerton, C.T., Hugenholtz, P., and Tyson, G.W. (2015) CheckM: Assessing the quality of microbial genomes recovered from isolates, single cells, and metagenomes. Genome Research 25: 1043–1055.

Puente-Sánchez, F., Arce-Rodríguez, A., Oggerin, M., García-Villadangos, M., Moreno-Paz, M., Blanco, Y., et al. (2018) Viable cyanobacteria in the deep continental subsurface. Proceedings of the National Academy of Sciences of the United States of America 115: 10702–10707.

Raven, J. (2009) Contributions of anoxygenic and oxygenic phototrophy and chemolithotrophy to carbon and oxygen fluxes in aquatic environments. Aquatic Microbial Ecology 56: 177–192.

Reno, M.L., Held, N.L., Fields, C.J., Burke, P. v., and Whitaker, R.J. (2009) Biogeography of the Sulfolobus islandicus pan-genome. Proceedings of the National Academy of Sciences of the United States of America 106: 8605–8610.

Rodriguez-R, L.M. and Konstantinidis, K.T. (2017) Bypassing Cultivation To Identify Bacterial Species Culture-independent genomic approaches identify credibly distinct clusters, avoid cultivation bias, and provide true insights into microbial species.

Roy, C., Rameez, M.J., Haldar, P.K., Peketi, A., Mondal, N., Bakshi, U., et al. (2020) Microbiome and ecology of a hot spring-microbialite system on the Trans-Himalayan Plateau. Sci Rep 10:.

Sánchez-Baracaldo, P. (2015) Origin of marine planktonic cyanobacteria. Sci Rep 5: 17418.

Saxena, R., Dhakan, D.B., Mittal, P., Waiker, P., Chowdhury, A., Ghatak, A., and Sharma, V.K. (2017) Metagenomic Analysis of Hot Springs in Central India Reveals Hydrocarbon Degrading Thermophiles and Pathways Essential for Survival in Extreme Environments. Front Microbiol 7: 2123.

Schirrmeister, B.E., Gugger, M., and Donoghue, P.C.J. (2015) Cyanobacteria and the Great Oxidation Event: evidence from genes and fossils. Palaeontology 58: 769–785.

Shi, T. and Falkowski, P.G. (2008) Genome evolution in cyanobacteria: The stable core and the variable shell. Proceedings of the National Academy of Sciences of the United States of America 105: 2510–2515.

Shih, P.M., Hemp, J., Ward, L.M., Matzke, N.J., and Fischer, W.W. (2017) Crown group Oxyphotobacteria postdate the rise of oxygen. Geobiology 15: 19–29.

Shih, P.M., Wu, D., Latifi, A., Axen, S.D., Fewer, D.P., Talla, E., et al. (2013) Improving the coverage of the cyanobacterial phylum using diversity-driven genome sequencing. Proceedings of the National Academy of Sciences of the United States of America 110: 1053–1058.

Soo, R.M., Hemp, J., and Hugenholtz, P. (2019) Evolution of photosynthesis and aerobic respiration in the cyanobacteria. Free Radical Biology and Medicine 140: 200–205.

Stamatakis, A. (2014) RAxML version 8: a tool for phylogenetic analysis and post-analysis of large phylogenies. Bioinformatics 30: 1312–1313.

Stolyar, S., Liu, Z., Thiel, V., Tomsho, L.P., Pinel, N., Nelson, W.C., et al. (2014) Genome sequence of the thermophilic cyanobacterium Thermosynechococcus sp. strain NK55a. Genome Announc 2: e01060–13.

Swanner, E.D., Mloszewska, A.M., Cirpka, O.A., Schoenberg, R., Konhauser, K.O., and Kappler, A. (2015) Modulation of oxygen production in Archaean oceans by episodes of Fe(II) toxicity. Nature Geoscience 8: 126–130.

Swanner, E.D., Wu, W., Hao, L., Wüstner, M.L., Obst, M., Moran, D.M., et al. (2015) Physiology, Fe(II) oxidation, and Fe mineral formation by a marine planktonic cyanobacterium grown under ferruginous conditions. Frontiers in Earth Science 3: 60.

Tang, J., Jiang, D., Luo, Y., Liang, Y., Li, L., Shah, Md.M.R., and Daroch, M. (2018) Potential new genera of cyanobacterial strains isolated from thermal springs of western Sichuan, China. Algal Research 31: 14–20.

Tomitani, A., Knoll, A.H., Cavanaugh, C.M., and Ohno, T. (2006) The evolutionary diversification of cyanobacteria: Molecular-phylogenetic and paleontological perspectives. Proceedings of the National Academy of Sciences of the United States of America 103: 5442–5447.

Toulza, E., Tagliabue, A., Blain, S., and Piganeau, G. (2012) Analysis of the Global Ocean Sampling (GOS) Project for Trends in Iron Uptake by Surface Ocean Microbes. PLoS ONE 7: e30931.

Ward, D.M., Ferris, M.J., Nold, S.C., and Bateson, M.M. (1998) A Natural View of Microbial Biodiversity within Hot Spring Cyanobacterial Mat Communities. Microbiol Mol Biol Rev 62: 1353–1370.

Ward, L.M., Fischer, W.W., and McGlynn, S.E. Candidatus Anthektikosiphon siderophilum OHK22, a New Member of the Chloroflexi Family Herpetosiphonaceae from Oku-okuhachikurou Onsen. Microbes Environ 35: 2020.

Ward, L.M., Idei, A., Nakagawa, M., Ueno, Y., Fischer, W.W., and McGlynn, S.E. (2019) Geochemical and Metagenomic Characterization of Jinata Onsen, a Proterozoic-Analog Hot Spring, Reveals Novel Microbial Diversity including Iron-Tolerant Phototrophs and Thermophilic Lithotrophs. Microbes and environments 34: 278–292.

Ward, L.M., Idei, A., Terajima, S., Kakegawa, T., Fischer, W.W., and McGlynn, S.E. (2017) Microbial diversity and iron oxidation at Okuoku-hachikurou Onsen, a Japanese hot spring analog of Precambrian iron formations. Geobiology 15: 817–835.

Ward, L.M. and Shih, P.M. (2019) The evolution and productivity of carbon fixation pathways in response to changes in oxygen concentration over geological time. Free Radical Biology and Medicine 140: 188–199.

Ward, L.M., Shih, P.M., and Fischer, W.W. (2018) MetaPOAP: presence or absence of metabolic pathways in metagenome-assembled genomes. Bioinformatics (Oxford, England) 34: 4284–4286.

Whitaker, R.J., Grogan, D.W., and Taylor, J.W. (2003) Geographic barriers isolate endemic populations of hyperthermophilic archaea. Science (80-) 301: 976–978.

Whitton, B.A. (1992) Diversity, ecology, and taxonomy of the cyanobacteria. In Photosynthetic prokaryotes. Springer, pp. 1–51.

Wu, Y., Zaiden, N., and Cao, B. (2018) The Core- and Pan-Genomic Analyses of the Genus Comamonas: From Environmental Adaptation to Potential Virulence. Frontiers in Microbiology 9: 3096.

Yeh, G.-H., Yang, T.F., Chen, J.-C., Chen, Y.-G., and Song, S.-R. (2005) Fluid geochemistry of mud volcanoes in Taiwan. Mud Volcanoes, Geodynamics and Seismicity Springer, Netherlands 227–237.

