## Supplemental Figure for "The *Thermosynechococcus* genus: wide environmental distribution, but a highly conserved genomic core"

### Supplementary Figure

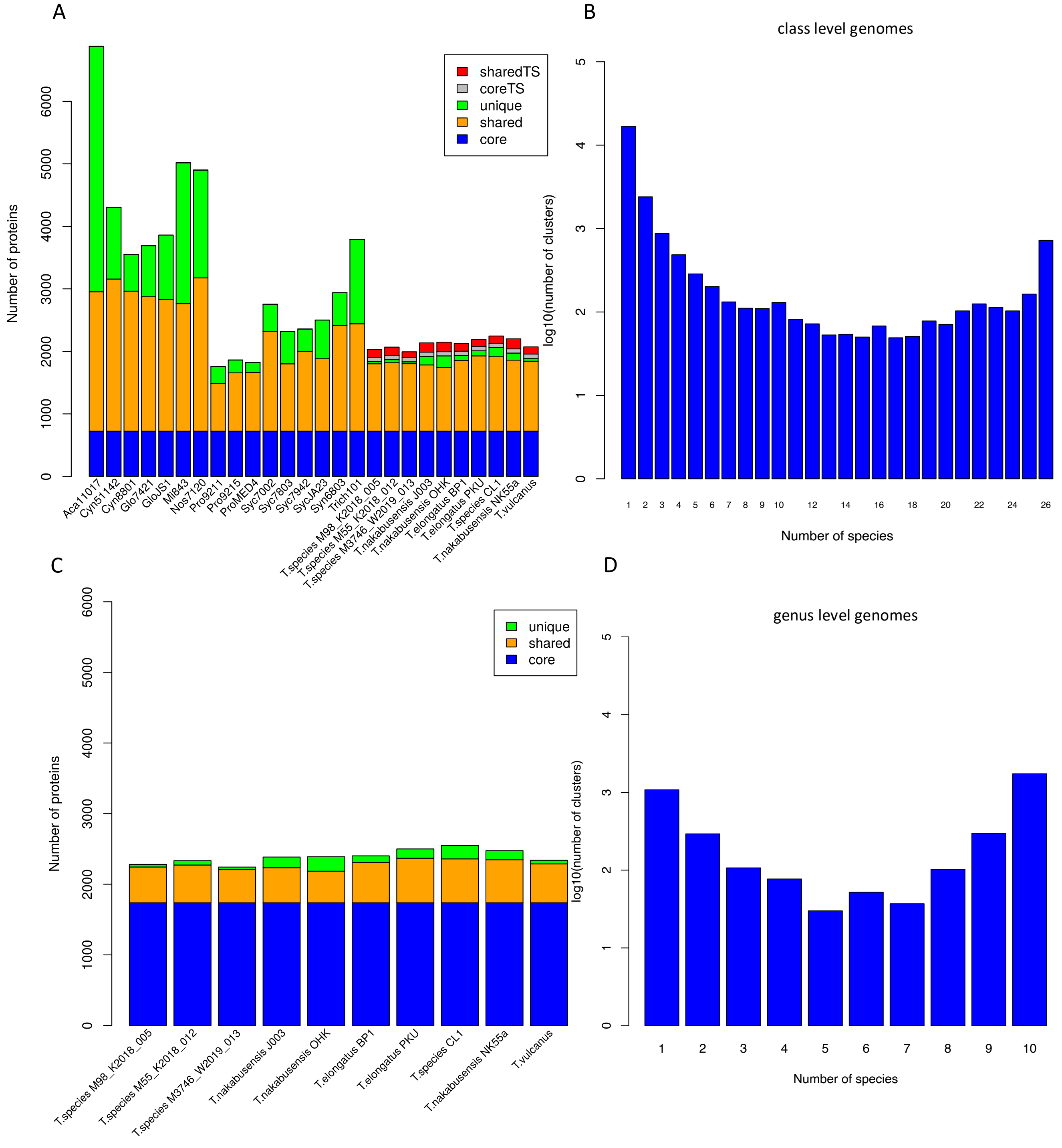


SI Figure S1: (A) Class level comparison of core- and pangenomes, (B) distribution of number of CLOGs with number of species at the class level, (C) genus level comparison of core- and pangenomes, (D) distribution of number of CLOGs with number of species at the genus level.
742 CLOGs are the core set of genes shared by all 23 cyanobacteria at the class level. The core at the class level seems to be stable when looking at the changes in the number of core CLOGs between 16 species (660 core CLOGs, Beck *et al* 2012), 77 species (621 core CLOGs, Beck *et al* 2018) and the analysis with 23 species here (742 core CLOGs).
